# A comparative study of ‘safe and just operating space’ for the south and south-east Asian countries

**DOI:** 10.1101/424200

**Authors:** Ajishnu Roy, Kousik Pramanick

**Affiliations:** Integrative Biology Research Unit, Department of Life Sciences, Presidency University, 86/1 College Street Kolkata – 700076

**Keywords:** Sustainable development goals, planetary boundaries, doughnut economy, safe and just operating space, national scale, South and Southeast Asia

## Abstract

The world is presently maintaining a delicate balance of continuing well-being and social development for the people through consumption of biophysical resources of nature without topping global average per capita availability. In this paper, we have framed a per capita top-down framework to survey national‘safe and just operating space’ (NSJOS) for the countries of south and southeast Asia to understand past variations and as a consequence, the present scenario. Amalgamating 27 indicators, all regarding Sustainable Development Goals (except – SDG 17), in consort with their respective environmental boundaries or desirable social development thresholds, this study explores into both biophysical (for ecological stress) and social development (for social deprivation) attributes of 19 countries of south and southeast Asia. This analysis shows, only 2 have remained either unchanged (political voice) or declining (social equity) among the 12 dimensions of social development in countries of this region. The remaining 10 dimensions of social development showing positive progress and will meet corresponding desired thresholds of United Nations Sustainable Development Goals 2015. All the 7 indicators showing tendencies of overconsumption of biophysical resources, that might be leading to exceeding per capita global average planetary boundaries in forthcoming future. However, ecological boundaries have remained protected to a decent degree so far for these countries. The challenge would be to maintain and increase the pace of social development and bringing it in equal strata of a global standard in future without depleting drivers of these, i.e. biophysical resources. National policy adaptations are crucial if these countries of south and southeast Asia desire to bring about adequacy in biophysical resources reserve whilst granting social equity in access and exploitation of these resources for the people towards the persistant social development in impending decades.

## Introduction

The era,‘Anthropocene’, symbolizes the scenario where anthropogenic drivers have become overriding forces of multifaceted changes in the Earth system (Steffen et al., 2007, 2011). This situation calls upon a comprehensive scientific understanding of biophysical (i.e. environmental) and socioeconomic interactions and concerned adaptive policy responses of local, regional, national to global decisions and development trajectories which might have unsustainable systemic multiscale adverse effects. The concept of sustainable development emerged for this context, that is “development that meets the needs of the present without compromising the ability of future generations to meet their own needs” (Brundtland Report, 1987). Agenda 21 (UNCED, Earth Summit, 1992) advised for sustainable development indicators (SDIs) to “provide solid bases for decision-making at all levels and to contribute to a self-regulating sustainability of integrated environment and development system”. 17 Sustainable Development Goals (SDGs) and 169 targets, ascended from Millennium Development Goals (MDGs) (Sachs, 2012) in 2015 by United Nations. These SDGs incorporate all 3 pillars of sustainable development, i.e. environmental goals (climate action, life below water, life on land etc.), economic goals (reduced inequalities, decent work and economic growth etc.) and social goals (zero hunger, no poverty, gender equality, peace and justice and strong institutions etc.). Two chief approaches have surfaced to quantified understanding of sustainability, (1) planetary boundaries (PBs) and (2) doughnut economy (DE), combining together forming safe and just operating space (SJS) framework. Rockström et al. introduced a new concept, ‘planetary boundaries’ framework, in 2009, to determine environment (i.e. ecological) thresholds and evaluate degree of consumption of nine biophysical resources that might prove to be hazardous to various Earth-system processes (climate change, rate of biodiversity loss, nitrogen and phosphorus cycles, stratospheric ozone depletion, ocean acidification, global freshwater use, change in land use, atmospheric aerosol loading and chemical pollution) (Rockström et al. 2009a, Rockström et al. 2009b). This framework (change in biosphere integrity, land-system change, the introduction of novel entities) was revised by Steffen et al. (2015). Dearing et al.’s (2014) defined ecological processes and control variables based on local environmental conditions of study locations, with a case study of two Chinese localities (Erhai lake-catchment, Yunnan province and Shucheng County, Anhui province, China). Nykvist et al. (2013) devised a methodology to determine national shares of four planetary boundaries (climate change, freshwater use, land-system change, and nitrogen), for 61 countries. Europe’s footprint was calculated by Hoff et al. (2014) with the PB framework. Dao et al. (2015) have analysed the sustainability of Switzerland through the PB framework. Nitrogen and phosphorus boundaries of Ethiopia and Finland was assessed by Kahiluoto et al. (2015). The planetary boundary of phosphorus was improved by Carpenter and Bennett (2011). Dao et al. (2018) have recently calculated the environmental limits of Switzerland accompanied with global limits, based on the PB framework. They have analysed PBs related to climate change, ocean acidification, nitrogen and phosphorus loss, land cover anthropisation and biodiversity loss. In a more recent report, Häyhä et al. (2018) have analysed the scope of safe operating space for countries in the European Union with the PB framework. They have incorporated six of nine indicators of PB (climate change, land system change, biogeochemical flows – nitrogen and phosphorus, freshwater use, biosphere integrity and novel entities). Parallelly, Raworth devised 11 dimensions of social foundation (water, income, education, resilience, voice, jobs, energy, social equity, gender equality, health and food), based on UNCSD (Rio+20, 2012) (Raworth, 2012). It was updated in 2017 to 12 dimensions (food, health, education, income and work, peace and justice, political voice, social equity, gender equality, housing, networks, energy, water) (Raworth, 2017a, 2017b). Cole et al. (2014) analysed the sustainable development of South Africa as ‘national barometer’ that included both planetary boundaries and doughnut economy frameworks with a culmination of both top-down and bottom-up approaches. They had modified few indicators from both frameworks (arable land use, air pollution and marine harvesting under the PB framework; health care, household goods, safety of SJS framework). O’Neill et al. (2018) have also downscaled these two frameworks towards national level analysis of 150 nations along with new indicators (e.g. eHANPP, ecological footprint, material footprint, life satisfaction, healthy life expectancy, nutrition, social support, democratic quality etc.). The recent Paris climate agreement advocates for an objective towards keeping global average temperature increase well below 2 degrees Celsius, if possible, to remain within 1.5 degrees Celsius for avoiding worst impacts. These objectives are needed to be achieved by the aggregate effects of national actions as Nationally Determined Contributions (NDCs). UN Convention on Biological Diversity (CBD) and UN Convention to Combat Desertification (UNCCD), both determine the necessary response at country level. These indicate to the significance of national-scale analysis. Our analysis measures the national-level performance of countries of south and southeast Asia on 28 dimensions, (both PB and SJS frameworks) and provides important outcomes concerning biophysical resource consumption and well-being for these countries. We have tried to understand how close these countries of south and southeast Asia have changed in their respective ‘safe’ ecological boundaries (i.e. national-level biophysical ceilings) (climate change, freshwater use, arable land use, nitrogen use, phosphorus use, ecological and material footprint) and how much of the population are socially deprived i.e. below ‘just’ social floor (i.e. national-level social foundations) (education, energy, food, gender equality, health, housing, income and work, networks, peace and justice, political voice, social equity, water and sanitation). This multinational analysis can yield a comparative overview of national-scale scenario so that every nation can comprehend which and where to focus enabling to meet UN SDG 2015 criteria.

## Data and Method

### a. Biophysical Indicators

We have used Rockström et al.’s (2009b) and Steffen et al.’s (2015) approach of planetary boundaries framework, with adjustment for all of the indicators and boundaries to fit national scale. Five indicators have been used from updated planetary boundaries framework of Steffen et al. (2015) (viz. climate change, nitrogen flow, phosphorus flow, land-system change and freshwater use) and two indicators from O’Neill et al.’s (2018) (ecological and material footprint).

#### Climate change

According to Rockström et al. (2009b), climate change boundary is based on global ‘atmospheric carbon dioxide concentration (parts per million by volume)’ and ‘change in radiative forcing i.e. energy imbalance at top-of-atmosphere (W m^-2^)’. For this dimension, Cole et al. (2014) used ‘annual direct CO_2_ emissions (Mt CO_2_)’ and O’Neill et al. (2018) used annual per capita CO_2_ emission (t CO_2_). We have measured this with greenhouse gas emission per capita per year. As per Emissions Gap Report (UNEP, November 2017), ‘emissions of all greenhouse gases should not exceed 42 GtCO_2_-e in 2030 if the 2 □ target is to be attained with higher than 66 per cent chance.’ This 42 GtCO_2_-e is divided with the global population to get per global capita scale boundary of 5.75 tCO_2_-e year^-1^ (2014).

#### Freshwater use

The Planetary boundary of freshwater use is the maximum withdrawal of 4000 km^3^ y^-1^ blue water from rivers, lakes, reservoirs, and renewable groundwater stores (Rockström et al., 2009b). Though Steffen et al. (2015) and O’Neill et al. (2018) followed this estimate, ‘annual consumption of available freshwater resources (Mm^3^ per year)’ has been used to measure this planetary boundary dimension by Cole et al. (2014). 4000 km^3^ y^-1^ water is divided with the global population to get global average per capita scale boundary of 574.86 km^3^ y^-1^ (2010).

#### Arable land use

The planetary boundary of land use is less than 15% of global ice-free land cover converted to cropland per year (which is 1995 Mha) (Rockström et al., 2009b). Steffen et al. (2015) have used ‘area of forested land as % of original forest cover’ and advised to maintain of 75% of global original forest cover, as a minimum (for tropical, temperate and boreal 85%, 50% and 85%, resp.). ‘Rain-fed arable land converted to cropland (%)’ has been used by Cole et al. (2014). O’Neill et al. (2018) have used ‘embodied human appropriation of net primary productivity (eHANPP)’ (ton Carbon per capita per year). For our analysis, we have divided 1995 Mha with the global population to get global average per capita scale land use boundary of 0.27ha year^-1^ (2015).

#### Nitrogen use

This boundary was measured with ‘amount of N_2_ removed from the atmosphere for human use (millions of tonnes y^-1^)’ (which was 35 million tonnes y^-1^) (Rockström et al., 2009b). According to Steffen et al. (2015), from industrial and intentional biological fixation, the planetary boundary of global N flow is 62 Tg N per year. O’Neill et al. (2018) have also used Steffen et al.’s (2015) method for their analysis. Cole et al. (2014) have used ‘nitrogen application rate of maize production (kg N ha^-1^)’. We have divided 62 Tg N y^-1^ with the global population to get the global average per capita scale boundary of 8.4kg N year^-1^ (2015).

#### Phosphorus use

This boundary was measured in terms of ‘quantity of phosphorus flowing into the ocean (millions of tonnes y^-1^)’ (i.e. global boundary of 11 million tonnes y^-1^) (Rockström et al., 2009b). However, according to Steffen et al. (2015), the planetary boundary of global phosphorus flow (mined and applied to erodible or agricultural soils) is 6.2 Tg N y^-1^. O’Neill et al. (2018) followed Steffen et al.’s (2015) method for this. Cole et al. (2014) used ‘total phosphorus concentration in dams (mg/L)’ as an indicator. We have divided 6.2 Tg N y^-1^ with the global population to get the global average per capita scale boundary of 0.84kg P year^-1^ (2015).

#### Ecological footprint (EF)

Ecological footprint is an indicator to measure how much biologically productive land and sea area a population requires to produce the biotic resources it consumes as well as absorb the CO_2_ emissions it generates, using prevailing technology and resource management practices (Borucke et al., 2013). It has been first used in the context of the planetary boundaries’ framework by O’Neill et al. (2018). According to Global Footprint Network (GFN), the world has 12 billion ha biologically productive land and sea area. We have divided 12 billion ha with the global population to get the global average per capita scale boundary of 1.66gha year^-1^ (2013).

#### Material footprint (MF)

Material footprint (also called Raw Material Consumption, RMC) is used to measure the amount of used material extraction (minerals, fossil fuels, and biomass) associated with the final demand for goods and services, irrespective of the location of the extraction (Wiedmann et al., 2015). It includes the embodied raw materials related to trade (both imports and exports) and thus, a suitable consumption-based measure. Global material footprint has been estimated at 70 Gt y^-1^ (i.e. 10.5 ton per capita in the year 2008, by Wiedmann et al. 2015), and it was capped to 8 ton per capita as a sustainable level, has been suggested by Dittrich et al. (2012). Global material extraction should not exceed ~50 Gt y^-1^, based on the material used in 2000 (50.8 Gt) (Dittrich et al., 2012). We have divided 50 Gt y^-1^ with the global population to get the global average per capita scale boundary of 7.18t year^-1^ (2010).

These dimensions (with respective indicators, boundaries, year, number of countries and data source) are explained in Table 1.

**Table 1:**
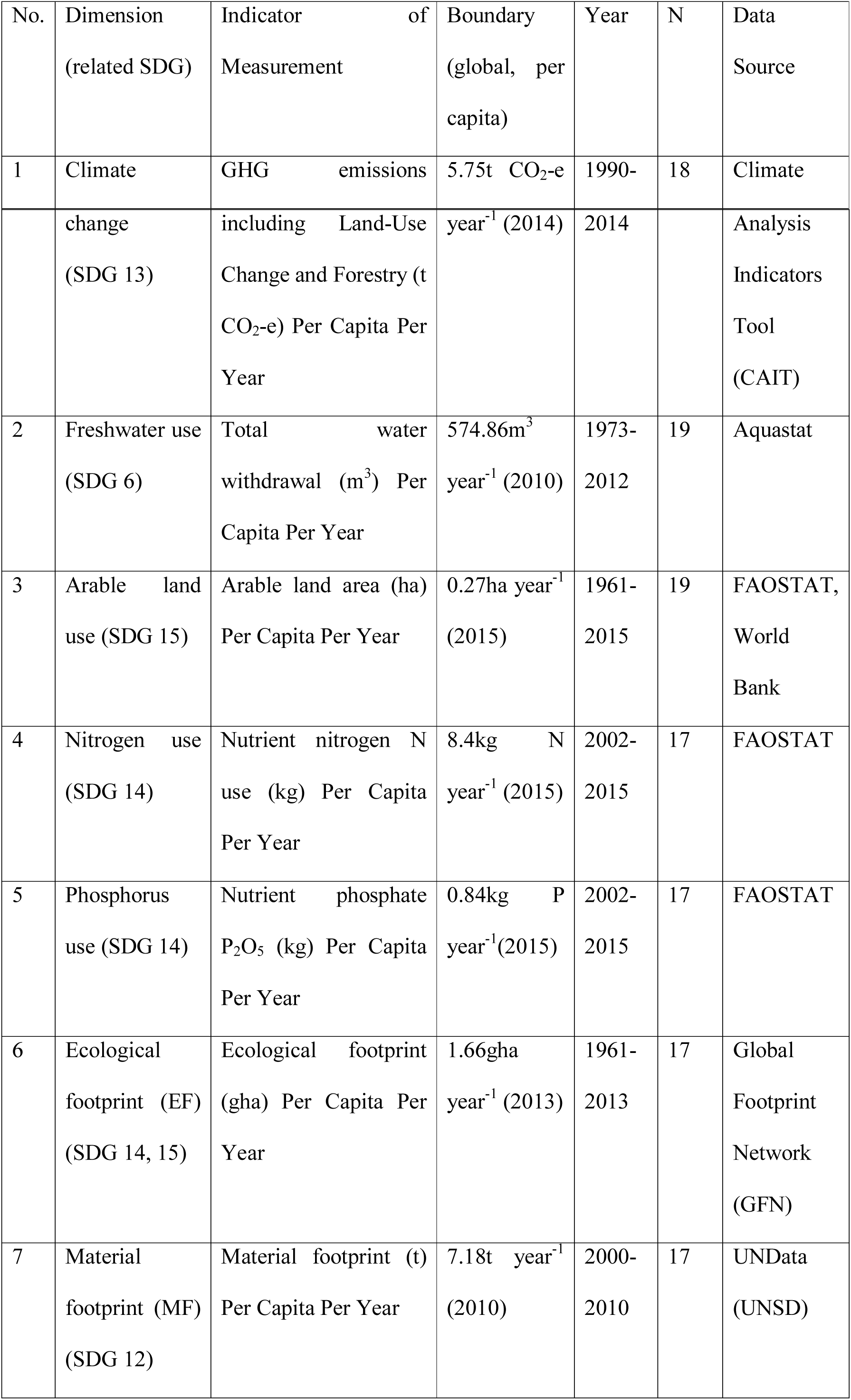
Dimensions and indicators of the ecological ceiling, related to safe operating space (SOS) of the Planetary boundaries concept.

### b. Social development Indicators

We followed Raworth’s (2017a) framework consisting of 12 dimensions related to social deprivation.

#### Education

SDG 4 aims at ensuring inclusive and equitable quality education and promotion of lifelong learning opportunities for all. 3 indicators have been selected from primary, secondary and adult education (literacy rate of adults, children of primary school age remained in school and secondary school enrollment) to reflect achievements and outcomes across diverse population age groups. Children out of school are the percentage of primary-school-age children who are not enrolled in primary or secondary school. Children in the official primary age group that are in pre-primary education should be considered out of school. The adult literacy rate is the percentage of people ages 15 and above who can both read and write with understanding a short simple statement about their everyday life. Gross enrollment ratio is the ratio of total enrollment, regardless of age, to the population of the age group that officially corresponds to the level of education shown. Secondary education completes the provision of basic education that began at the primary level and aims at laying the foundations for lifelong learning and human development, by offering more subject- or skill-oriented instruction using more specialized teachers. We have used the data from the World Bank’s World Development Indicators (WDI). A threshold of 10% or less has been chosen for children out of the school of primary school age and 90% or more was chosen for secondary school enrollment and adult literacy rate.

#### Energy

SDG 7 aims to ensure access to affordable, reliable, sustainable and clean energy for all. About 1 billion people currently do not have access to electricity. 3 billion people rely on polluting fuel (like – fuelwood, charcoal, crop residue, animal dung, dry leaves) to cook food, which in turn resulting in 4 million premature deaths per year, mostly among women and children, that are due to household air pollution (SDG 7 tracking report, 2018). Our assessment of deprivations in access to energy includes both electricity and the quality of (clean) cooking facilities. We have measured energy using two indicators, (1) ‘access to electricity (% of populations)’ and (2) ‘access to clean fuels and technologies for cooking (% of the population)’, obtained from the World Bank’s WDI. Access to electricity is the percentage of the population with access to electricity. Access to clean fuels and technologies for cooking is the proportion of total population primarily using clean cooking fuels and technologies for cooking. Under WHO guidelines, kerosene is excluded from clean cooking fuels. The threshold for energy was set at 90% or more for both indicators.

#### Food

The target of SDG 2 is ending hunger, achieving food security and improved nutrition for all. We measured social development related to food using two indicators, (1) ‘average calorific intake of food & drink (kcal/capita/day)’ and (2) ‘prevalence of undernourishment (% of the population)’. Daily caloric supply is defined as the average per capita caloric availability. This indicates the caloric availability delivered to households but does not necessarily indicate the number of calories actually consumed (food may be wasted at the consumer level). The physiological requirements for an average adult remain between 2100 and 2900 kcal per day (for average men and women with moderate physical activity). This calorific requirement range exceeds for individuals associated with heavy manual labour or athletic activity (Smil, 2000). We have followed O’Neill et al. (2018) and used 2700 kcal or more per capita day y^-1^ as a threshold. Population below minimum level of dietary energy consumption (also referred to as prevalence of undernourishment) shows the percentage of the population whose food intake is insufficient to meet dietary energy requirements continuously.

#### Gender Equality

The focus of SDG 5 is achieving gender equality via empowering all women. It would be ideal to assess the extent of gender inequality to understand women and men’s roles and status in political and economic life. We measured this using one indicator - ‘proportion of seats held by women in national parliaments (%)’ from the World Bank’s World Development Indicators. Women in parliaments are the percentage of parliamentary seats in a single or lower chamber held by women. The indicator value is calculated such that if women held exactly half of all parliamentary seats (i.e. 50%), that should be non-biased to both genders. Thus, achieving 50% seats in parliament has been taken as the desired threshold.

#### Health

Ensuring healthy lives and promoting well-being for all at all ages is the focus of SDG 3. We have used two indicators to assess shortfalls in access to health care in India: (1) ‘life expectancy at birth, total (years)’ and (2) ‘mortality rate, <5 years (per 1,000 live births)’ from the World Bank’s WDI. Life expectancy at birth indicates the number of years a newborn infant would live if prevailing patterns of mortality at the time of its birth were to stay the same throughout its life. Seventy years or more life expectancy at birth is selected here as a desirable threshold (Human Development Report, UNDP, 2015). The under-five mortality rate is the probability per 1,000 that a newborn baby will die before reaching age five, if subject to age-specific mortality rates of the specified year. The international target for all countries to reduce under-five years age mortality to at least as low as 25 per 1,000 live births by 2030 (WHO, 2015). Thus, 25 or less per 1000 live births has been set as the desired threshold here.

#### Housing

SDG 11 focus on making cities and human settlements inclusive, safe, resilient and sustainable. We have measured it with ‘population living in slums (% of urban population)’ from WDI (World Bank). Population living in slums is the proportion of the urban population living in slum households. A slum household is defined as a group of individuals living under the same roof lacking one or more of the following conditions: lack of access to improved drinking water, lack of access to improved sanitation, overcrowding (>3 persons per room) and dwellings made of non-durable material. We have set the threshold at 10% or less of urban population living in slums.

#### Income and Work

SDG 1 focus on ending poverty in all its forms everywhere. Promoting sustained, inclusive, sustainable economic growth full and productive employment and decent work for all is the goal of SDG 8. We have used (1) ‘poverty headcount ratio at $1.90 a day (2011 PPP) (% of the population)’ to measure income and (2) ‘unemployment, youth total (% of total labour force, 15-24 years)’ for work, both from WDI (World Bank). Poverty headcount ratio at $1.90 a day is the percentage of the population living on less than $1.90 a day at 2011 international prices. Though the goal is having 100% of the population living above the $1.90 a day line, we have used 95% as a threshold value in this analysis. Youth unemployment refers to the share of the labour force ages 15-24 years without work, though available for and seek employment (International Labour Organization, ILO estimation). We have used 94% or more people are employed (i.e. 6% or less unemployed people below this line) is the desired threshold.

#### Networks

SDG 9 focus on building resilient infrastructure, promoting inclusive and sustainable industrialization and fostering innovation. Under this goal, target 9.c. focus on significantly increasing access to information and communications technology and strive to provide universal and affordable access to the Internet in the least developed nations. The network has been measured with ‘individuals using the internet (% of the population)’ provided by WDI of the World Bank. Internet users are individuals who have used the Internet (from any location) in the last 3 months. The Internet can be used via a computer, mobile phone, personal digital assistant, games machine, digital TV etc. We have used 90% or more of the population have access to the internet as the desired threshold.

#### Peace & Justice

UN SDG 16 focus on promoting peaceful and inclusive societies for sustainable development, provide access to justice for all and build effective, accountable and inclusive institutions at all levels. We used two indicators (1) corruption perceptions index (CPI) for justice, (provided by Transparency International) and (2) ‘intentional homicides (per 100,000 people)’ (from WDI) for peace. Corruption perceptions index scores countries according to how corrupt their public sector is perceived to be, i.e. highly corrupt to very clean [scale: 0 to 10 (up to 2011) and 0 to 100 (2012 onwards)]. We have set the desired threshold of 5 or less (up to 2011) and 50 or less (2012 onwards). Intentional homicides are estimates of unlawful homicides purposely inflicted as a result of domestic disputes, interpersonal violence, violent conflicts over land resources, intergang violence over turf or control, and predatory violence and killing by armed groups. It does not include all intentional killing. Individuals or small groups usually commit homicide, whereas killing in armed conflict is usually committed by fairly cohesive groups of up to several hundred members and is thus usually excluded. We have set the threshold at 10 or fewer homicide deaths per 100,000 population per year.

#### Political Voice

Under SDG 16, target 16.7 aims for ensuring responsive, inclusive, participatory and representative decision-making at all levels. We have measured political voice using voice & accountability index (VAI), provided as a component of the World Bank’s World Governance Indicators (WGI). Voice and accountability capture perceptions of the extent to which a country’s citizens are able to participate in selecting their government, as well as freedom of expression, freedom of association, and a free media. This index is scored on a scale of 0 to 1 (i.e. very poor performance to very high performance). The threshold is set at 0.5 or less on this indicator.

#### Social Equity

SDG 10 focus on reducing inequality within and among the countries. The shortfall of social equity is measured with national income inequalities. We have measured social equity using the Gini coefficient of per capita expenditure, provided by the UNU-WIDER, World Income Inequality Database 3.4 (WIID 3.4) (Jan, 2017). Gini index measures the extent to which the distribution of income (or, in some cases, consumption expenditure) among individuals or households within an economy deviates from a perfectly equal distribution. A Gini index of 0 represents perfect equality, while an index of 100 implies perfect inequality. The threshold was chosen of 70 of 0-100 scale of Gini index of 0.30.

#### Water & Sanitation

SDG 6 focus on ensuring availability and sustainable management of water and sanitation for all. Deprivations in access to water and sanitation services are assessed on the basis of two widely used indicators, (1) ‘improved sanitation facilities (% of the population with access)’ and (2) ‘improved water source (% of the population with access)’ from the World Bank’s WDI. Access to improved sanitation facilities refers to the percentage of the population using improved sanitation facilities. Improved sanitation facilities are likely to ensure hygienic separation of human excreta from human contact. They include flush/pour flush (to the piped sewer system, septic tank, pit latrine), ventilated improved pit (VIP) latrine, pit latrine with slab, and composting toilet. Although it is preferable that 100% of the population should have access to improved sanitation facilities, we have chosen a threshold of 90% for this indicator. Access to an improved water source refers to the percentage of the population using an improved drinking water source. The improved drinking water source includes piped water on premises (piped household water connection located inside the user’s dwelling, plot or yard), and other improved drinking water sources (public taps or standpipes, tube wells or boreholes, protected dug wells, protected springs, and rainwater collection). We have set the threshold for this indicator as 90% or more people have access (i.e. 10 or fewer people do not have access) to an improved water source.

These dimensions (with respective indicators, boundaries, year, number of countries and data source) are explained in Table 2.

**Table 2.**
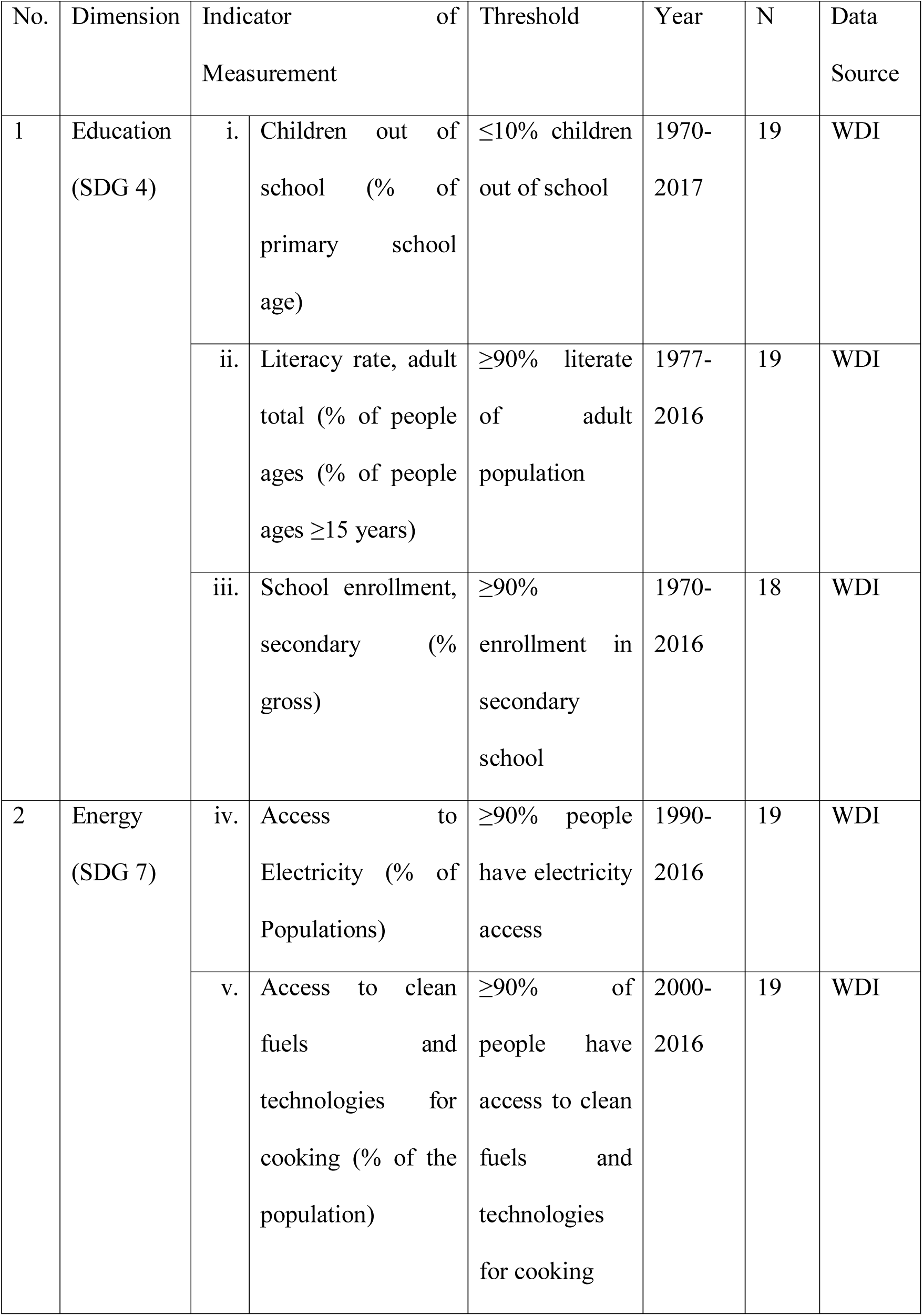
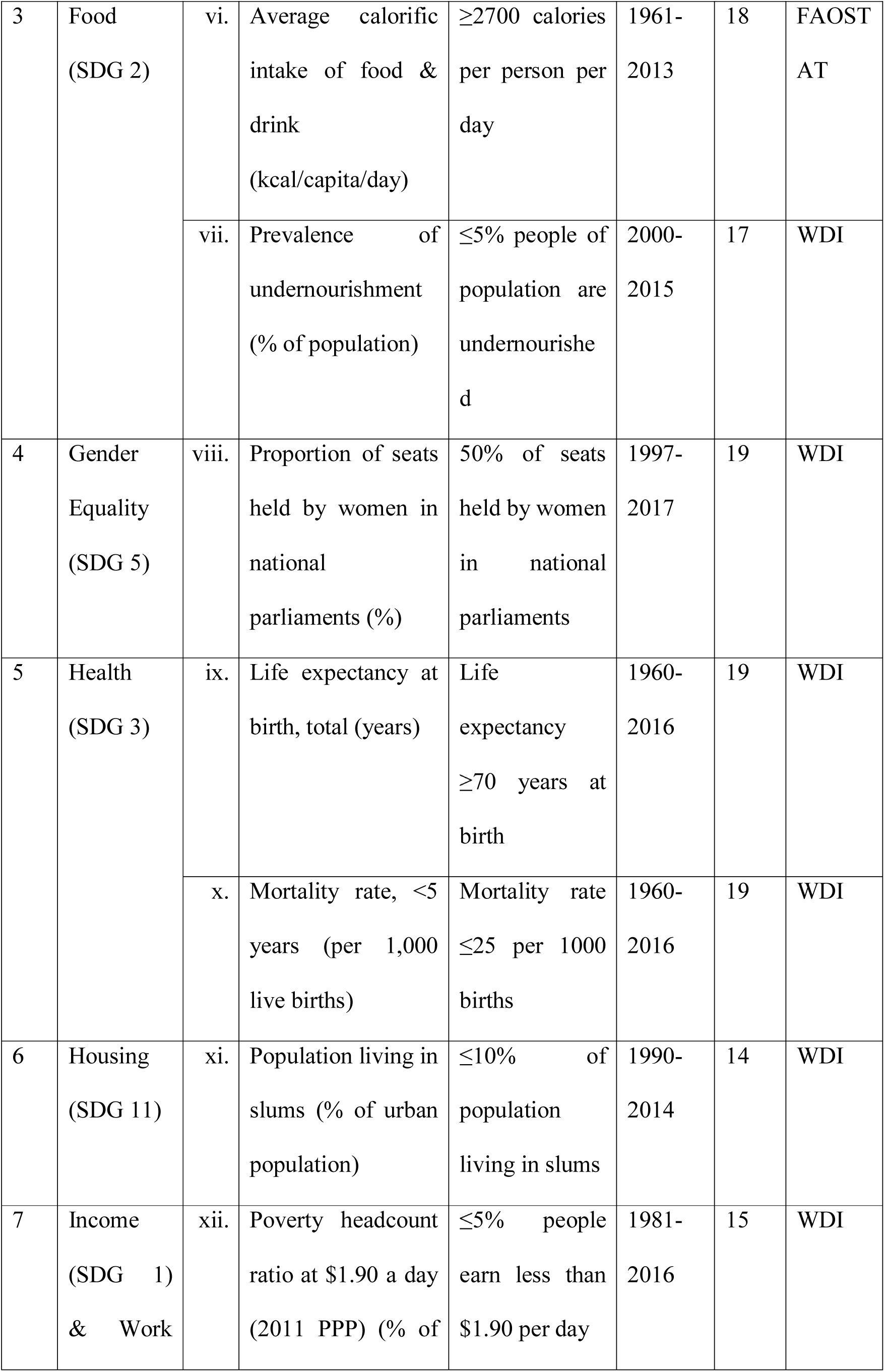
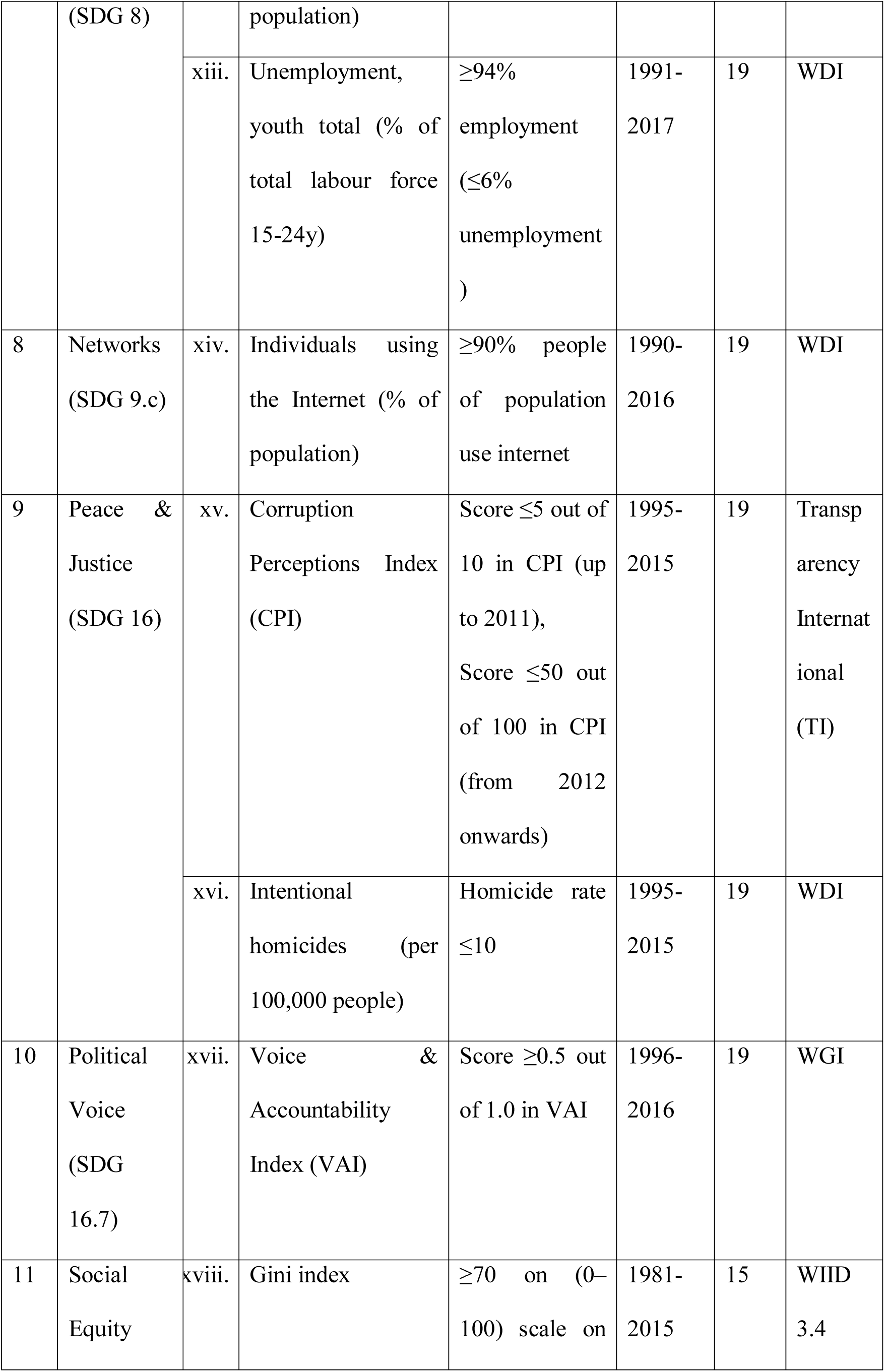
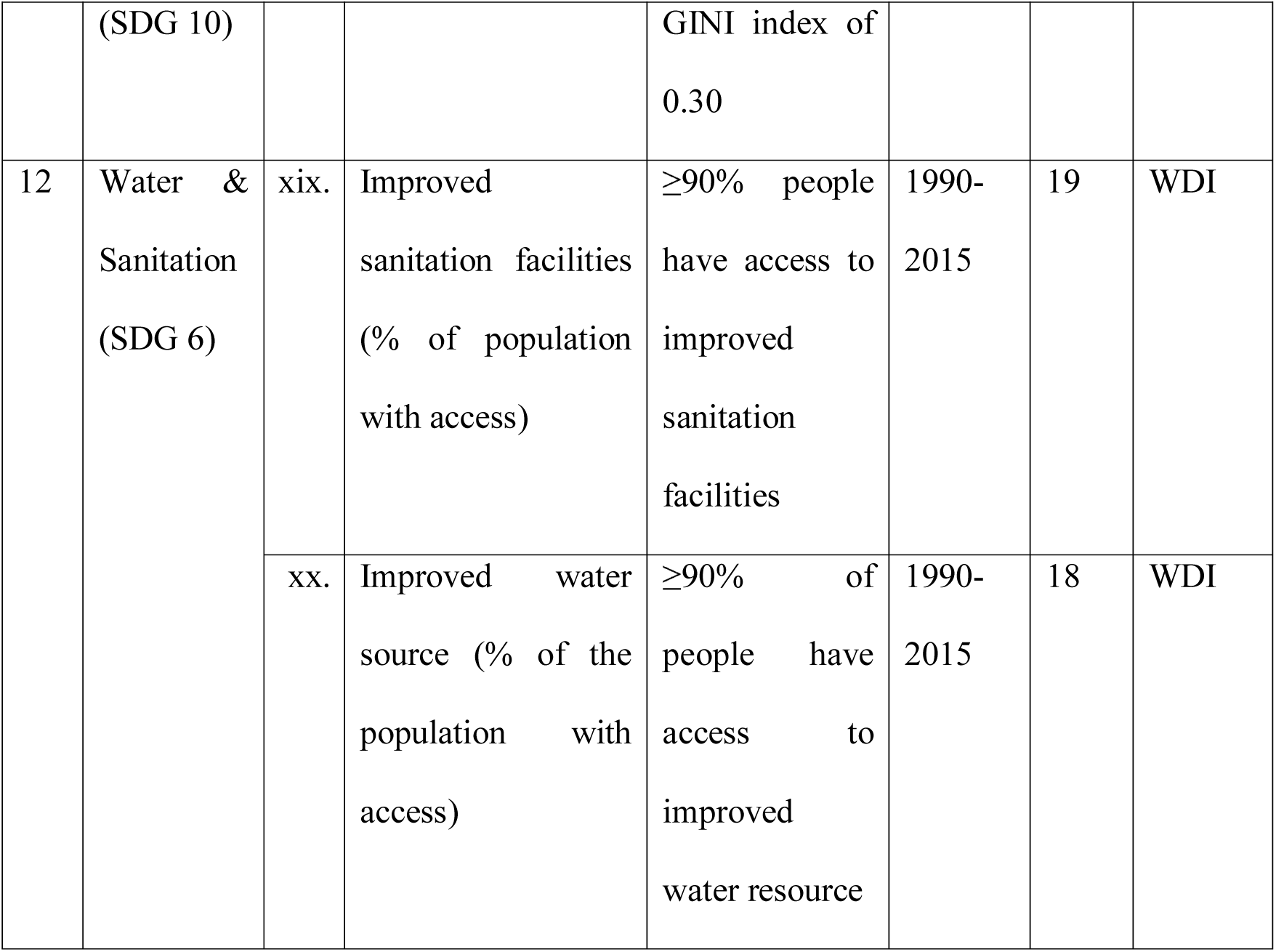
Dimensions and indicators of the social foundation, related to safe and Just space (SJS) of Doughnut economy concept.

| No. | Dimension | Indicator of Measurement | Threshold | Year | N | Data Source |
| --- | --- | --- | --- | --- | --- | --- |
| 1 | Education (SDG 4) | i. Children out of school (% of primary school age) | ≤10% children out of school | 1970-2017 | 19 | WDI |
|  |  | ii. Literacy rate, adult total (% of people ages (% of people ages ≥15 years) | ≥90% literate of adult population | 1977-2016 | 19 | WDI |
|  |  | iii. School enrollment, secondary (% gross) | ≥90% enrollment in secondary school | 1970-2016 | 18 | WDI |
| 2 | Energy (SDG 7) | iv. Access to Electricity (% of Populations) | ≥90% people have electricity access | 1990-2016 | 19 | WDI |
|  |  | v. Access to clean fuels and technologies for cooking (% of the population) | ≥90% of people have access to clean fuels and technologies for cooking | 2000-2016 | 19 | WDI |

|  |  |  |  |  |  |  |  |
| --- | --- | --- | --- | --- | --- | --- | --- |
| 3 | Food<br>(SDG 2) | vi. | Average calorific intake of food & drink<br>(kcal/capita/day) | ≥2700 calories per person per day | 1961-2013 | 18 | FAOST<br>AT |
|  |  | vii. | Prevalence of undernourishment<br>(% of population) | ≤5% people of population are undernourished | 2000-2015 | 17 | WDI |
| 4 | Gender Equality<br>(SDG 5) | viii. | Proportion of seats held by women in national parliaments (%) | 50% of seats held by women in national parliaments | 1997-2017 | 19 | WDI |
| 5 | Health<br>(SDG 3) | ix. | Life expectancy at birth, total (years) | Life expectancy ≥70 years at birth | 1960-2016 | 19 | WDI |
|  |  | x. | Mortality rate, <5 years (per 1,000 live births) | Mortality rate ≤25 per 1000 births | 1960-2016 | 19 | WDI |
| 6 | Housing<br>(SDG 11) | xi. | Population living in slums (% of urban population) | ≤10% of population living in slums | 1990-2014 | 14 | WDI |
| 7 | Income<br>(SDG 1)<br>& Work | xii. | Poverty headcount ratio at \$1.90 a day (2011 PPP) (% of | ≤5% people earn less than \$1.90 per day | 1981-2016 | 15 | WDI |
|  | (SDG 8) |  | population) |  |  |  |  |
|  |  | xiii. | Unemployment, youth total (% of total labour force 15-24y) | ≥94% employment (≤6% unemployment) | 1991-2017 | 19 | WDI |
| 8 | Networks (SDG 9.c) | xiv. | Individuals using the Internet (% of population) | ≥90% people of population use internet | 1990-2016 | 19 | WDI |
| 9 | Peace & Justice (SDG 16) | xv. | Corruption Perceptions Index (CPI) | Score ≤5 out of 10 in CPI (up to 2011), Score ≤50 out of 100 in CPI (from 2012 onwards) | 1995-2015 | 19 | Transparency International (TI) |
|  |  | xvi. | Intentional homicides (per 100,000 people) | Homicide rate ≤10 | 1995-2015 | 19 | WDI |
| 10 | Political Voice (SDG 16.7) | xvii. | Voice & Accountability Index (VAI) | Score ≥0.5 out of 1.0 in VAI | 1996-2016 | 19 | WGI |
| 11 | Social Equity | xviii. | Gini index | ≥70 on (0–100) scale on | 1981-2015 | 15 | WIID 3.4 |
|  | (SDG 10) |  |  | GINI index of<br>0.30 |  |  |  |
| 12 | Water &<br>Sanitation<br>(SDG 6) | xix. | Improved<br>sanitation facilities<br>(% of population<br>with access) | ≥90% people<br>have access to<br>improved<br>sanitation<br>facilities | 1990-2015 | 19 | WDI |
|  |  | xx. | Improved water<br>source (% of the<br>population with<br>access) | ≥90% of<br>people have<br>access to<br>improved<br>water resource | 1990-2015 | 18 | WDI |

## Results

### a. Biophysical Indicators

#### Climate change

Global average per capita GHG emission had crossed climate change PB in 2005 and stayed ahead since then. Among 18 countries, 4 has already crossed climate change PB (Brunei, Indonesia, Singapore and Malaysia, in highest to lowest order of per capita GHG emission) and 5 are on the way to cross Climate change PB within a few years (Thailand, Laos, Myanmar, Maldives and Cambodia, in highest to lowest order of GHG emission). Only one country, Bhutan has shown negative GHG emission per capita, but its GHG absorbing capacity has been decreasing over the years. Remaining 8 countries are within the safe limits of climate change PB. Also, among 18 countries, 5 have shown the positive scenario of decreasing per capita GHG emission (Cambodia, Malaysia, Nepal, Philippines and Singapore) and the rest, 13 countries along with the world, are showing increasing per capita GHG emission with time.

#### Freshwater use

Among 19 countries, 9 have already crossed per capita freshwater use PB (Afghanistan, India, Myanmar, Pakistan, Philippines, Sri Lanka, Thailand, Timor-Leste and Vietnam) and 3 are on its way to cross within a few years (Bhutan, Indonesia and Laos). Remaining 7 are within the safe limits of freshwater use PB. Here, we would like to add that scarcity of data in the Aquastat database acts as a hindrance to a more comprehensive understanding of freshwater use, especially for these countries of south and southeast Asia.

#### Land use

World average per capita available arable land use is presently within the safe limits of arable land use PB. However, as the global population continues to increase, both arable land use PB and global average per capita arable land use decrease. Problem is evident when it is clear from data that the gap between these two are closing in fast and if this increasing rate of population growth remains uninterrupted for a few years, World will cross the safe limits of arable land use PB. Among the 19 countries, though none has crossed the safe limits of arable land use PB, 5 have come very close to cross the safe limits within a few years (Afghanistan, Cambodia, Laos, Myanmar and Thailand). Only one country, Sri Lanka, has shown increased per capita arable land use as opposed to all other remaining 18 countries.

#### Nitrogen use

Much before 2002, World had crossed nitrogen use PB and the gap between global average per capita nitrogen use and safe limits of nitrogen use PB is continuously increasing over the years. Among the 17 countries, 9 has already crossed nitrogen use PB (Thailand, Vietnam, Pakistan, India, Indonesia, Afghanistan, Sri Lanka, Malaysia and Bangladesh, in decreasing order of per capita nitrogen use) and another 4 are close to crossing safe limits of N use PB (Cambodia, Nepal, Philippines and Bhutan, in decreasing order of per capita nitrogen use). Remaining 4 countries are well within the safe limits of nitrogen use PB. It is clear from the data that even the country with the highest value of nitrogen use is almost 50% less than the global average value. From this, we can infer that, from the perspective of per capita nitrogen use, these countries of south and southeast Asia are not responsible for such high global average values of nitrogen use and as a consequence, crossing of the safe limits of nitrogen use PB.

#### Phosphorus use

The world had crossed phosphorus use PB well before 2002 and the gap between safe limits of phosphorus use PB and global average per capita phosphorus use is continuously increasing with time. Among the 17 countries, 12 has already crossed phosphorus use PB (Cambodia, Philippines, Bhutan, Nepal, Sri Lanka, Indonesia, Bangladesh, Thailand, India, Pakistan, Vietnam and Malaysia, in increasing order of per capita phosphorus use) and 2 are closing in to cross the phosphorus use PB (Brunei and Maldives). Remaining 3 are well within the safe limits of phosphorus use PB.

#### Ecological footprint

Among 17 countries of south and southeast Asia, 6 have already crossed ecological footprint PB (Vietnam, Laos, Thailand, Malaysia, Bhutan and Singapore, in increasing order of per capita ecological footprint) and 7 others are close to crossing the ecological footprint PB (Cambodia, India, Indonesia, Myanmar, Nepal, Philippines and Sri Lanka). Remaining 4 are well within the safe limits of ecological footprint PB.

#### Material footprint

Among 17 countries, 5 has already crossed material footprint PB (Bhutan, Thailand, Maldives, Malaysia and Singapore, in increasing order of per capita material footprint) and another 6 are about to cross material footprint PB within a few years (India, Cambodia, Philippines, Laos, Indonesia and Viet Nam, in increasing order of per capita material footprint). Remaining 6 are well within the safe limits of material footprint PB.

### b. Social development Indicators

#### Education

The world reached the desired threshold (10% or less) in ‘children out of the school of primary school age’ in 2007 and improving since. Among 19 countries, though 5 have not reached the threshold (Afghanistan, Bhutan, Pakistan, Thailand and Timor-Leste), remaining all 14 countries have already reached it. In recent few years, for 3 countries (Indonesia, Thailand and Timor-Leste), the distance from threshold is increased.

The world is yet to reach the desired threshold (90% or more) in ‘adult literacy rate’, however, the condition is improving continuously. Among 19 countries, 9 have already reached the threshold (Brunei, Indonesia, Malaysia, Maldives, Philippines, Singapore, Sri Lanka, Thailand and Vietnam) and 4 others are close to reaching it (Bangladesh, Cambodia, India and Myanmar).

The world is yet to reach the desired threshold (90% or more) in ‘secondary school enrolment rate’, however, the condition is improving with time. Among 18 countries, only 2 have reached the threshold (Brunei and Sri Lanka) and 8 others are closing in (Bhutan, India, Indonesia, Malaysia, Nepal, Philippines, Thailand and Timor-Leste).

#### Energy

Though the condition is improving, World is far from reaching the desired threshold (90% or more) in ‘access to clean fuels and technologies for cooking’. Among 19 countries, only 4 have reached the threshold (Brunei, Malaysia, Maldives and Singapore) and 3 others are close to reaching it (Indonesia, Thailand and Vietnam). However, the remaining 12 are far away from reaching the threshold.

The world is yet to achieve the target threshold (90% or more people) in ‘access to electricity’. Among 19 countries of south and southeast Asia, 12 have reached the threshold (Bhutan, Brunei, Indonesia, Malaysia, Maldives, Nepal, Pakistan, Philippines, Singapore, Sri Lanka, Thailand and Vietnam) and 4 are very close among the remaining (Afghanistan, Bangladesh, India, and Laos).

#### Food

The world has already reached the desired threshold (2700 kcal or more day^-1^) for ‘average calorific intake of food & drink’ in 1998. Among 17 countries, only 6 have reached the threshold (Brunei, Indonesia, Malaysia, Maldives, Vietnam and Thailand) and 4 (Myanmar, Nepal, Philippines and Sri Lanka) are very close to reaching it among the remaining 11.

The world has not yet reached the desired threshold (5% or less) for ‘prevalence of undernourishment’. Among 17 countries, only 2 have reached the threshold (Brunei and Malaysia) and 4 are very close among the remaining (Indonesia, Maldives, Nepal and Thailand).

#### Gender Equality

The world has reached only half of the desired threshold (50%) for ‘proportion of seats held by women in national parliaments. Although among the 19 countries, none has reached threshold, 6 countries have achieved almost equal or better than the global average (Afghanistan, Laos, Nepal, Philippines, Timor-Leste and Vietnam) and 9 are far away from reaching it in near future (Bhutan, Brunei, India, Indonesia, Malaysia, Maldives, Myanmar, Sri Lanka and Thailand).

#### Health

The world has already reached the desired threshold (70 years or more) for ‘life expectancy at birth’ in 2008 and continuously improving since. Among 19 countries of south and southeast Asia, 10 have reached the threshold (Bangladesh, Bhutan, Brunei, Malaysia, Maldives, Nepal, Singapore, Sri Lanka, Thailand and Vietnam). All the remaining 9 countries are presently very close and have the potential to reach the threshold within a few years.

Though the condition is improving, the world is yet to reach the desired threshold (25 or less per 1000 live births) in ‘mortality rate of fewer than 5 years’ olds. Among 19 countries, only 7 have reached the threshold (Brunei, Malaysia, Maldives, Singapore, Sri Lanka, Thailand and Vietnam). 6 countries among the remaining 12 are very close to reaching the threshold (Bangladesh, Bhutan, Cambodia, Indonesia, Nepal and Philippines).

#### Housing

The world is far for achieving the threshold (10% or less) in ‘population living in slums in urban areas’ and the condition is not improving at all. Among the 14 countries, none has reached the threshold but, 7 are close to it (India, Laos, Pakistan, Singapore, Sri Lanka, Timor-Leste and Vietnam).

#### Income and Work

Though the condition is improving, the world is yet to reach the desired threshold (5% or less) in ‘poverty headcount ratio at $1.90 a day (2011 PPP)’. Among 15 countries, 5 have reached the threshold (Bhutan, Malaysia, Sri Lanka, Thailand and Vietnam) and 5 of the remaining are close to it (Indonesia, Maldives, Myanmar, Pakistan and Philippines).

The world is yet to reach the desired threshold (6% or less) in ‘youth unemployment, 15-24y’ and also, the condition is deteriorating globally since 2008. Among 19 countries, only 6 have reached the threshold (Cambodia, Laos, Myanmar, Nepal, Singapore and Thailand) and 3 of the rest are close to it (Pakistan, Philippines and Vietnam).

#### Networks

Though the scenario is improving rapidly, the world has just reached only half of the desired threshold (90% or more) in ‘individuals using the internet’. Among 19 countries, only a single country, Brunei, has achieved the threshold and 2 are close to it (Malaysia and Singapore). All others are either near global average or even more distant from the threshold.

#### Peace and Justice

Among 19 countries, 3 have yet to reach the threshold (5 or less, up to 2011 and 50 or less from 2012 onwards) in ‘corruption perceptions index’ and all the remaining 16 have achieved the threshold. In general, the CPI score is increasing i.e. the level of corruption is getting higher gradually.

The world has reached the desired threshold (10 or less) for ‘intentional homicides (per 100,000 people)’ before 2012. Among 19 countries, all have reached the threshold. However, the condition is towards a fall for 4 countries with time (Afghanistan, Laos, Pakistan and Philippines).

#### Political voice

Among 19 countries, none have achieved the threshold [more than 0.5 in -2.5 (weak) to +2.5 (strong) scale] of ‘voice and accountability index’. However, 4 countries are closer to the threshold than the rest (India, Indonesia, Philippines and Timor-Leste).

#### Social inequality

Among 15 countries, none have achieved the threshold [70 on (0–100) scale] of the Gini index. However, 2 countries are closer to the threshold than the rest (Philippines and Myanmar).

#### Water and Sanitation

Though the global scenario of sanitation is improving, the world is yet to reach the desired threshold (90% or more) in ‘improved sanitation facilities. Among 18 countries, 5 have reached the threshold (Malaysia, Maldives, Singapore, Sri Lanka and Thailand) and 4 are closing in (Laos, Myanmar, Philippines and Vietnam).

The world has already reached the desired threshold (90% or more) of ‘improved water source’ in 2013 and improving since then. Among 18 countries, 11 have reached the threshold (Bhutan, India, Malaysia, Maldives, Nepal, Pakistan, Philippines, Singapore, Sri Lanka, Thailand and Vietnam) and 3 are getting close to it (Bangladesh, Indonesia and Myanmar).

The cumulative scenario of 27 indicators related to safe and just operating space for the countries of south and southeast Asia of the present time (as per latest available data) in contrast with the past (the 2000s) are represented in Table 3. All of the biophysical resource consumption indicators are deteriorating. Among the 12 domains of social development, only 2 have remained either unchanged (political voice) or declining (social equity). In all of the remaining 10 domains, the overall scenario for these countries showing positive changes. This means development is taking place at the cost of overconsumption of the biophysical resources. Ranking of countries based on this SJS score is given in Supplementary Table 1.

**Table 3:**
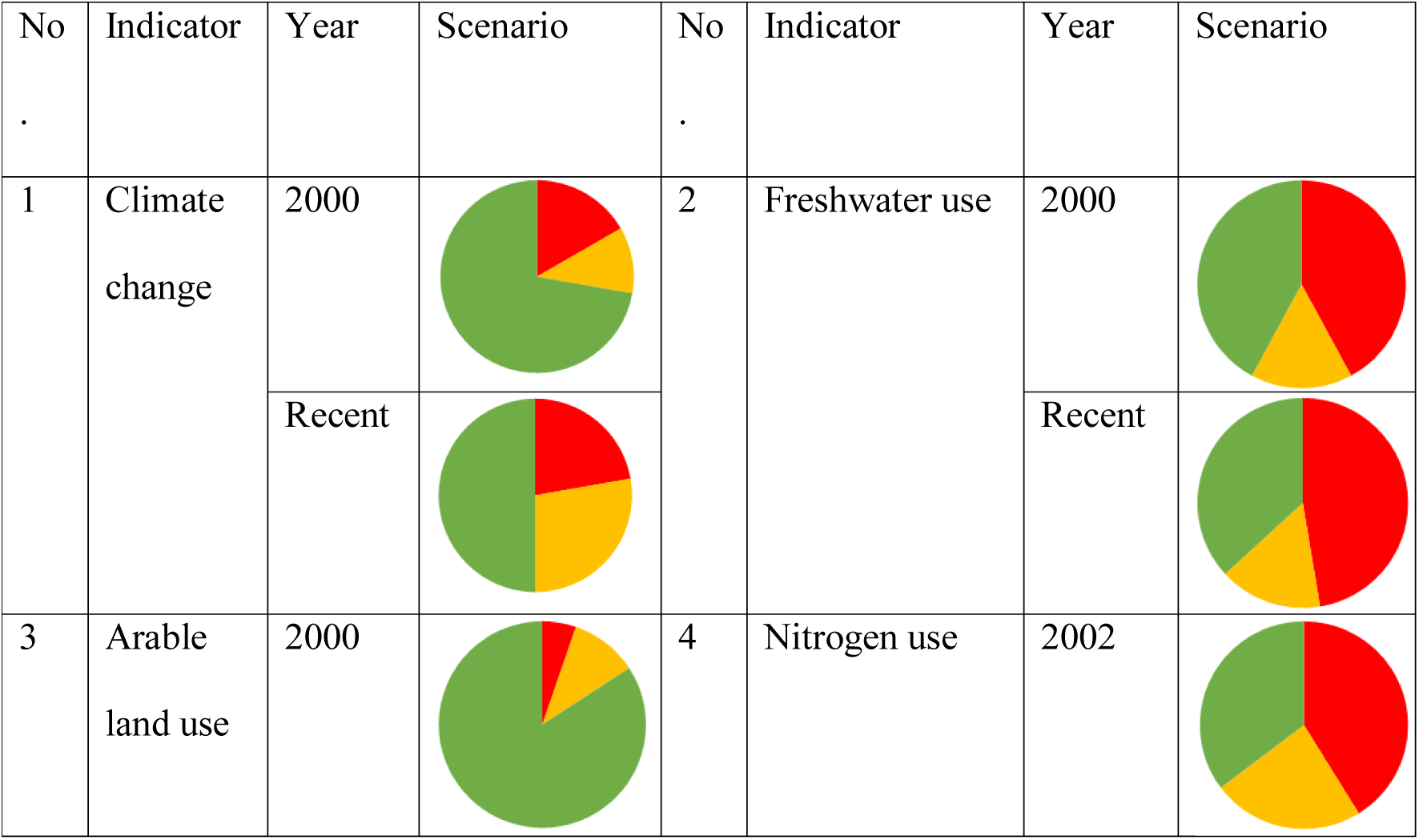
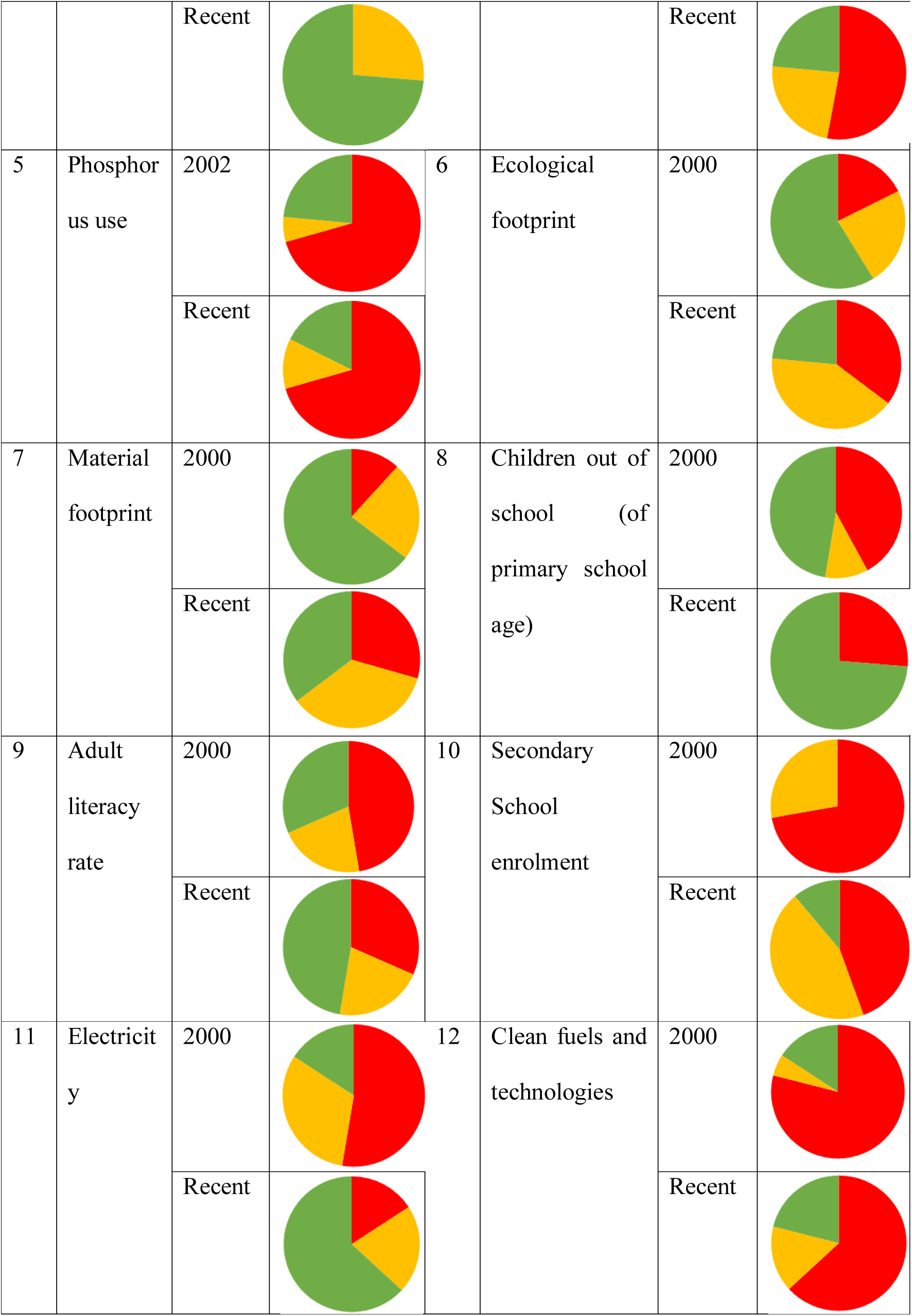
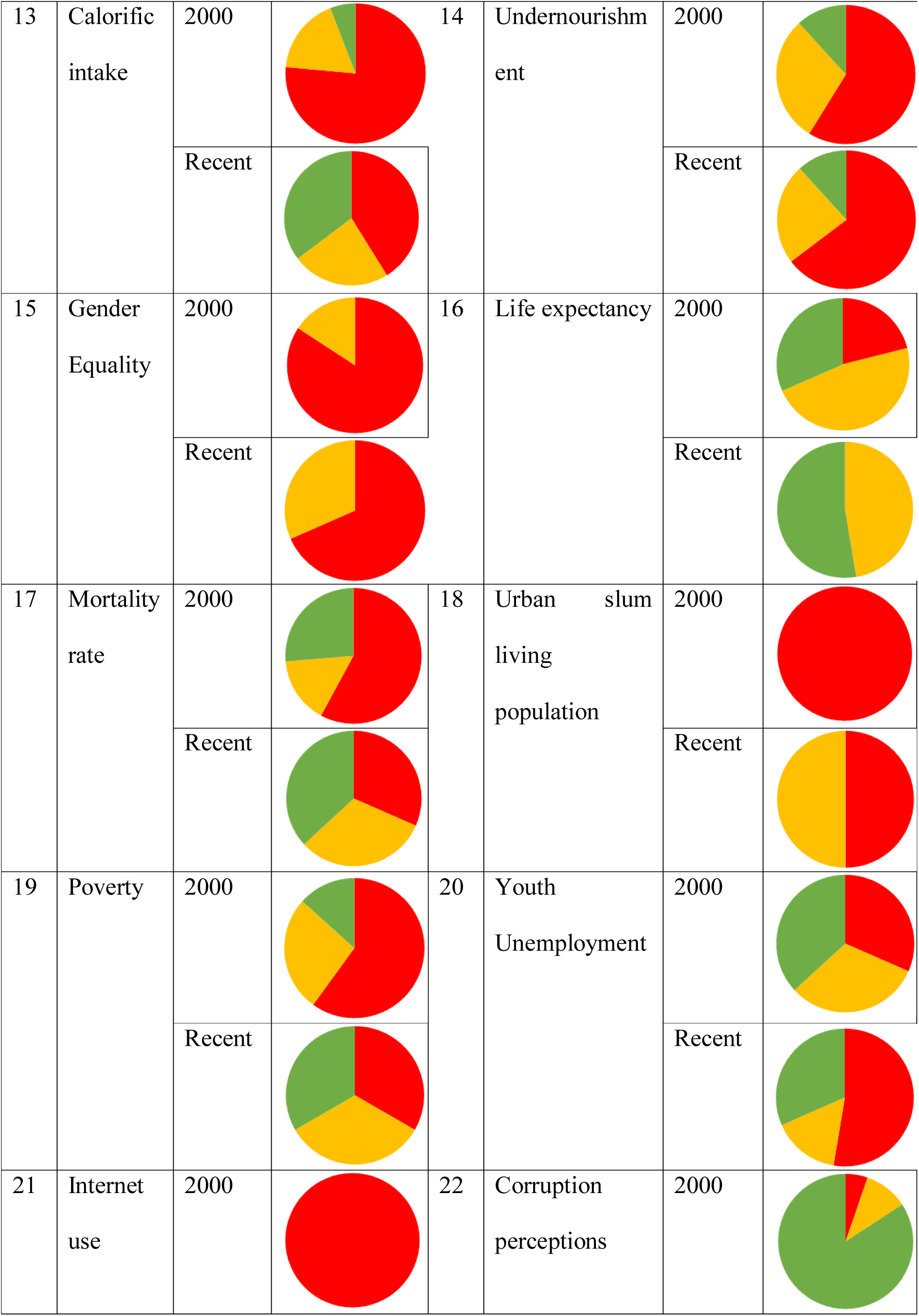
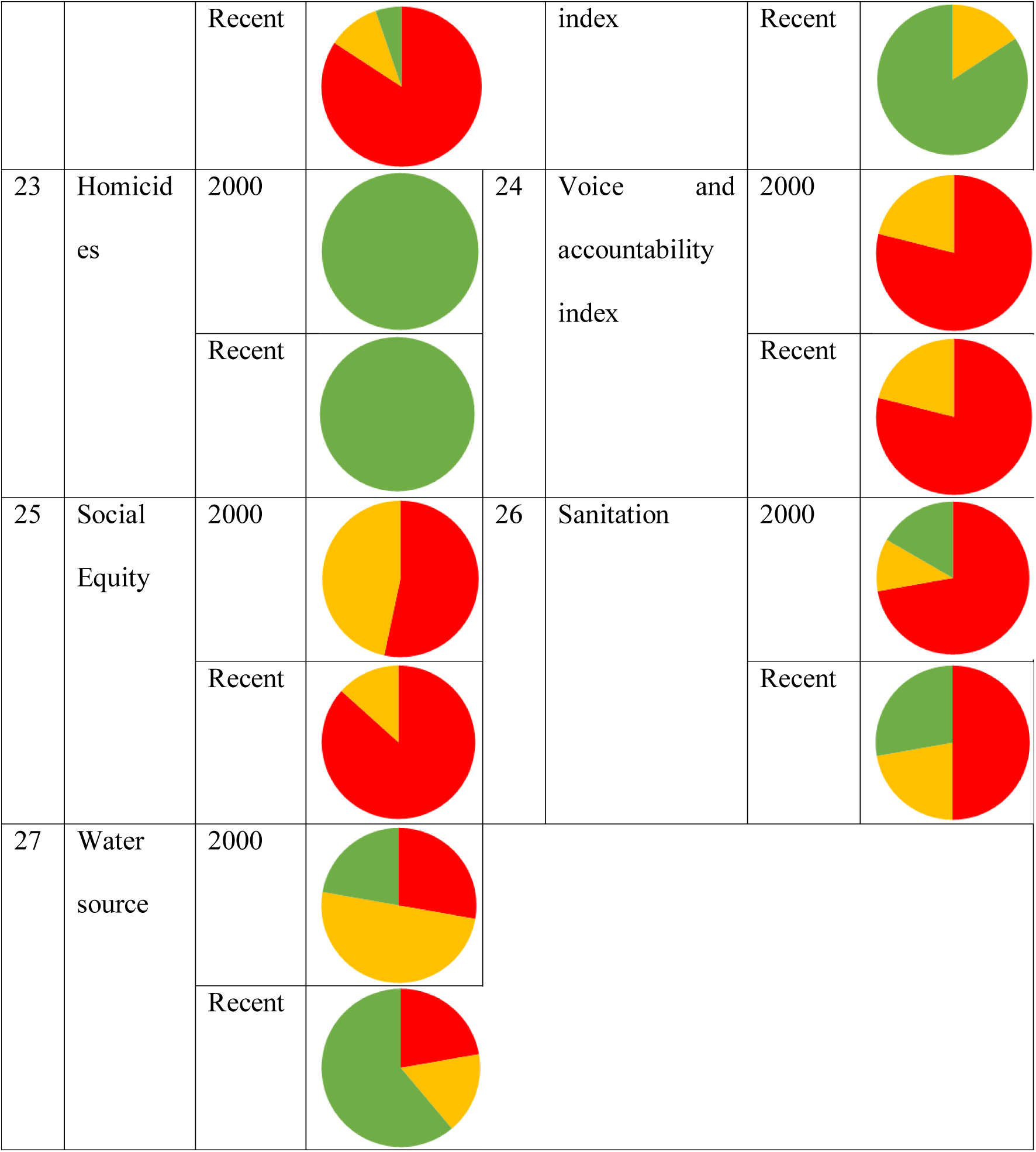
Trends of indicators related to safe and just operating space (SJS) for countries of south and southeast Asia.

For biophysical indicators (1-7), green indicates the proportion of countries within the safe limits of PB, red indicates the proportion of countries that have crossed biophysical boundaries and yellow indicates the proportion of countries that are close to crossing the biophysical boundaries. For social development indicators (8-27), green indicates the proportion of countries who have achieved the sustainable development goals set by United Nations, yellow indicates the proportion of countries close to reaching the targets and red indicates the proportion of countries far distant from the desired goals.

We get a safe and just space location graph (safe in the x-axis and just in y-axis) for countries of south and southeast Asia in Fig. 3. From this graph, it can be clearly understood that – (1) Only 2 countries are in a stage where, a higher degree of social development has been achieved at a cost of depleted biophysical resources, namely Malaysia and Thailand. (2) The only single country is in the zone of safe and just space is the Maldives. (3) Other 3 countries have shown desired progress by entering the zone of safe and just space, viz. Brunei, Sri Lanka and Vietnam. (4) However, most of the countries are in a zone of safe space i.e. they have not crossed most of the respective biophysical boundaries, but, also not significant progress has been achieved in social development. To sum up, most of the countries of south and southeast Asia belong to the zone of safe but not just operating space, neither in past (the 2000s) nor in present (as per most recently available data).

**Fig. 1.**
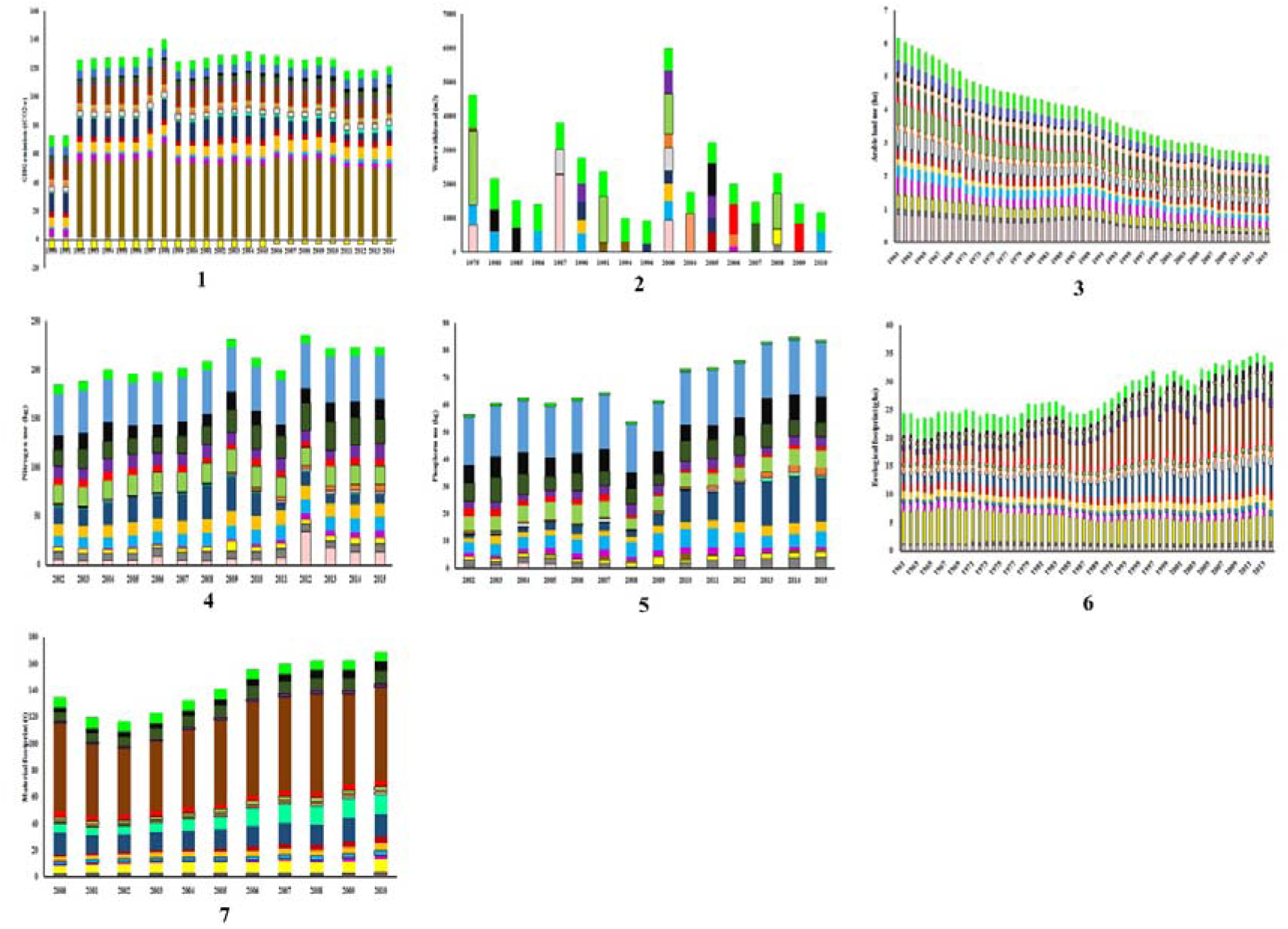
Trends in biophysical indicators related to Planetary boundaries for countries of South and Southeast Asia. Biophysical indicators are – (1) GHG emission, (2) water withdrawal, (3) arable land use, (4) nitrogen use, (5) phosphorus use, (6) ecological footprint and (7) material footprint. 19 countries considered here, are – Afghanistan (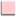), Bangladesh (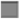), Bhutan (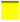), Brunei (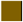), Cambodia (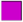), India (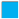), Indonesia (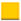), Lao PDR (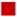), Malaysia (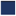), Maldives (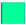, Myanmar (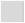), Nepal (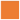), Pakistan (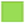), Philippines (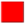), Singapore (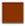), Sri Lanka (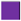), Thailand (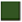), Timor-Leste (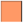) and Vietnam (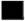). Global average values (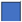) and respective planetary average per capita boundaries(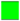) have also been shown.

**Fig. 2.**
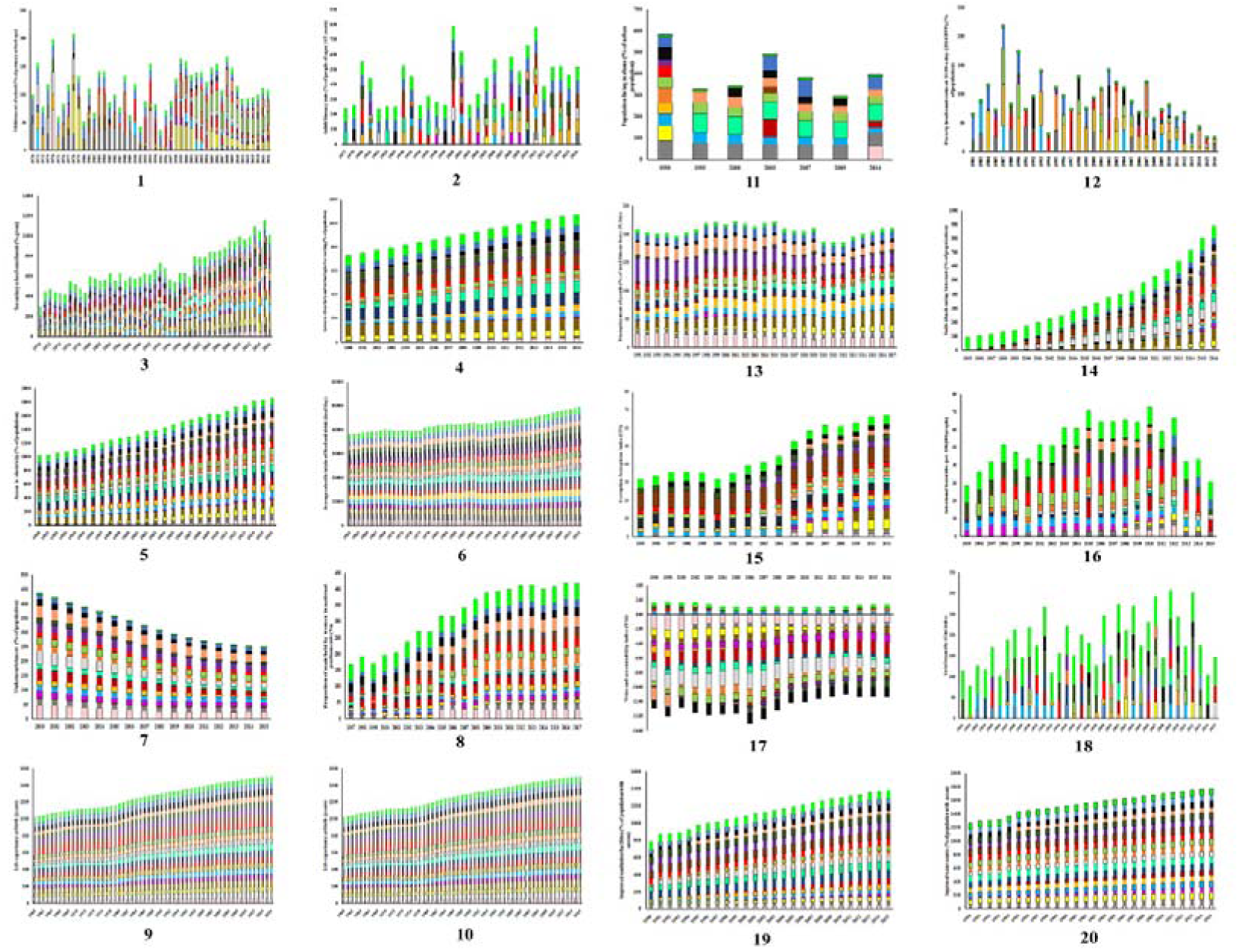
Trends in social development indicators related to safe and Just space (SJS) of Doughnut economy for countries of South and Southeast Asia. Indicators of social development are – (1) children out of school of primary school age, (2) adult literacy rate, (3) secondary school enrolment, (4) access to electricity, (5) access to clean fuels and technologies for cooking, (6) average calorific intake of food and drink, (7) undernourishment, (8) proportion of seats held by women in national parliaments, (9) life expectancy at birth, (10) mortality rate under 5 years, (11) urban population living in slums, (12) poverty headcount ratio at $1.90 a day, (13) youth unemployment, (14) individuals using the internet, (15) corruption perceptions index, (16) intentional homicides, (17) voice and accountability index, (18) Gini index, (19) improved sanitation facilities and (20) improved water source. Twelve dimensions of the social foundation are education 1-3, energy 4-5, food 6-7, gender equality 8, health 9-10, housing 11, income and work 12-13, networks 14, peace and justice 15-16, political voice 17, social equity 18, water and sanitation 19-20. 19 countries considered here, are – Afghanistan (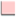), Bangladesh (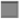), Bhutan (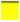), Brunei (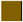), Cambodia (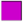), India (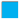), Indonesia (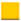), Lao PDR (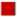), Malaysia (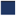), Maldives (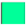, Myanmar (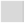), Nepal (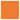), Pakistan (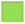), Philippines (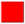), Singapore (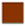), Sri Lanka (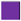), Thailand (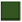), Timor-Leste (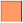) and Vietnam (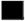). Global average values (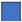) and respective United Nations sustainable development goal values (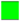) have also been shown.

**Fig. 3.**
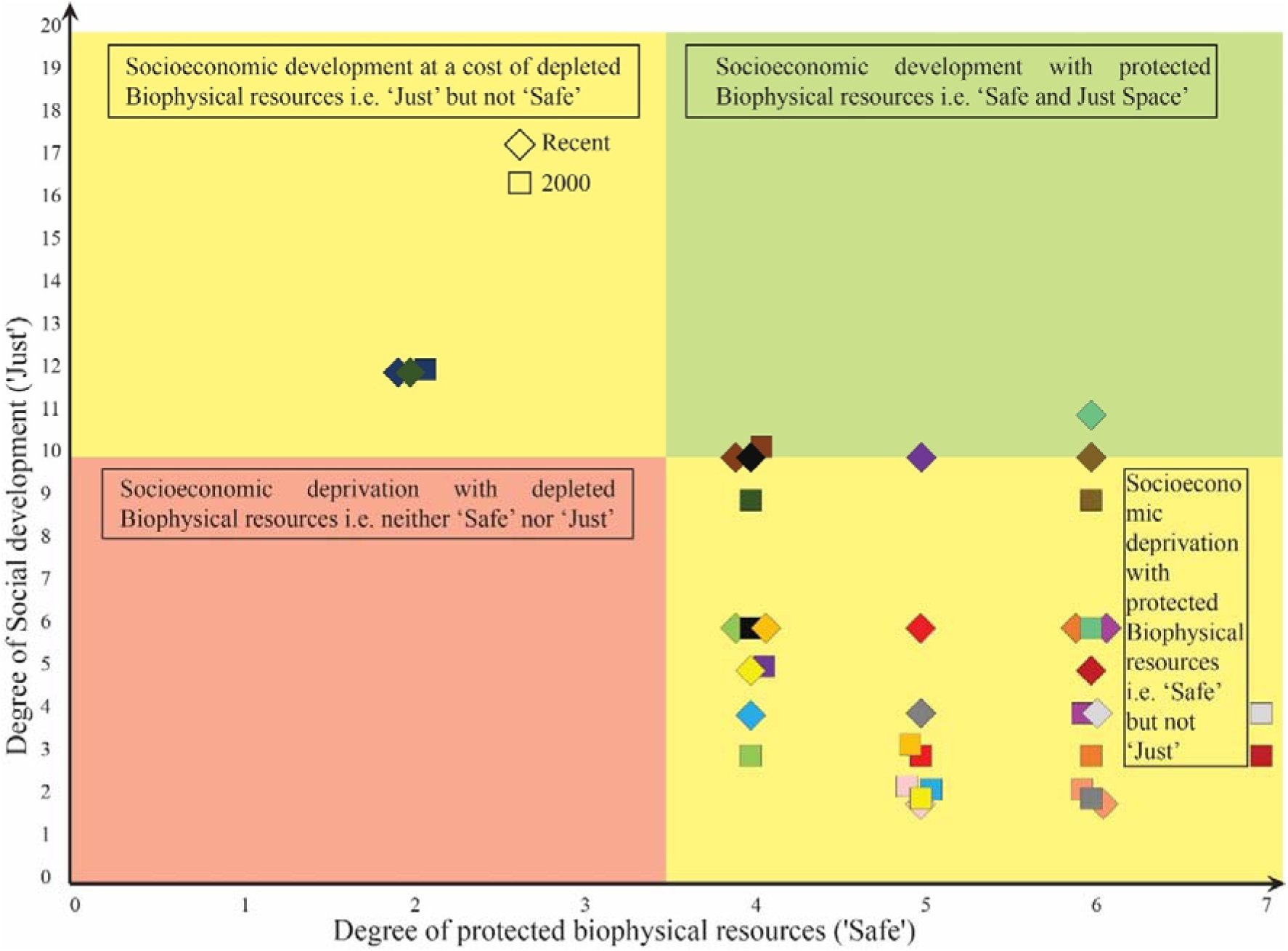
Presence of countries (N=19) of south and southeast Asia in past (2000, shown as rhombus) and present (as per recent available data, shown as square) on number of social development thresholds achieved (‘just’ space, 20 indicators) versus number of biophysical boundaries not crossed (‘safe’ space, 7 indicators). The location has been divided into 4 zones – I. neither safe or just zone (lower left), II. safe but not just zone (lower right), III. just but not safe zone (upper left) and IV. Safe and just zone (upper right). 19 countries considered here, are – Afghanistan (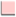), Bangladesh (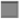), Bhutan (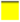), Brunei (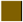), Cambodia (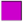), India (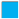), Indonesia (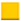), Lao PDR (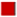), Malaysia (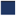), Maldives (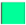, Myanmar (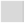), Nepal (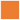), Pakistan (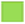), Philippines (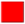, Singapore (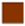), Sri Lanka (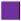), Thailand (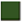), Timor-Leste (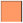) and Vietnam (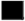).

## Discussion

There have been a lot of interdisciplinary studies for the last few years allied to SJS framework. DE framework for both Welsh and UK has been analyzed by Sayers and Trebeck (2015) and Sayers (2015). Chapron et al. (2017) advocated towards enforcement of environmental laws in form of tools to check anthropogenic impacts on the environment through playing under safe limits of planetary boundaries. The ‘biosphere integrity’ has created a lot of debate surrounding a suitable indicator for quantification (Samper, 2009; Running, 2012; Mace et al., 2014; Newbold et al., 2016). Likewise, for ‘freshwater use’ (Rockström and Karlberg, 2010; Bogardi et al., 2013; Gerten et al., 2013; Heistermann, 2017, Gleick, 2018) ‘introduction of novel entities’ (Sala and Goralczyk, 2013; Persson et al., 2013; Diamond et al., 2015; Villarrubia-Gómez et al., 2017) with their respective safe boundaries. There has also been some work to connect the planetary boundary with governance, along with policy implications (Bierman, 2012, Galaz et al., 2012a, 2012b; Reischl, 2012). A significant amount of work has also been done on establishing and applying PB framework at regional-scale (Dearing et al., 2014; Häyhä et al., 2016; Cole et al., 2017; McLaughlin, 2018). Some studies have been done to explore the connection of food system and nutrients with PB framework (Kahiluoto et al., 2014, 2015; Campbell et al., 2017; Conijn et al., 2018). A preliminary framework to apply the PB framework in the marine context has been done by Nash et al. (2017). Simultaneously, there have been criticisms of SJS framework (Montoya et al., 2018a, 2018b). Recently, Steffen et al. (2018) have proposed the scenario of ‘Hothouse Earth’ as a consequence of unchecked biophysical consumption if humanity continues to cross the safe limits of planetary boundaries and maintain that trajectory.

Two studies have incorporated sustainability of these countries of south and south-east Asia based on PB and SJS framework, Nykvist et al. (2013) and O’Neill et al (2018). Nykvist et al. (2013) have considered four planetary boundaries in their report, (1) climate change (tCO_2_ per capita y^1^), (2) nitrogen use (kg N per capita y^-1^), (3) freshwater use (m^3^ per capita y^-1^) and (4) land use (ha per capita). According to them, most of these countries did not cross per capita PB. One deficit in the first study is that it did not include correlated social dimensions (i.e. ‘just space’ framework or any other). O’Neill et al. (2018) used seven and eleven indicators for ‘safe’ and ‘just’ space analysis, respectively. According to them, although these countries have trans relatively fewer biophysical boundaries, they have also failed to achieve very few social thresholds in comparison with other developed countries of the world.

The principal aim of this study was to evaluate the sustainable development, inclusively, using SJS framework at the national level for countries of south and south-east Asia. We have tried to maintain the original concept and design of the two frameworks as much as possible while deriving results that are meaningful in national-scale for countries of south and southeast Asia.

There are some recommendations that we came up during this study: (1) There is an increasing necessity to establish and maintain the publicly available sub-national level database. (2) Data coverage period should be as long as possible. This might prove to be a good opportunity for data-poor countries to commence a competent accumulation of inclusive key data for addressing their national-to-global challenges. (3) Fitting indicator for analysing the progress for the original Steffen’s (2015) and Raworth’s (2017) framework should be cultivated. (4) Additional work is needed for an approach to recognize policymaking and implementation gaps of each nation for all of the indicators in the SJS framework. (5) The biophysical resource consumption (‘safe’ part) is not at yet most endangered for these countries. But social development (‘just’ part) should be need of the hour for these countries, which remaining under safe limits of biophysical resources. (6) We recommend that every nation (like – countries of south and southeast Asia) should act more preemptively and embrace policies in accordance with the recommendations of international authoritative organizations, like - UN, UNFCC, UNDP, UNEP etc. (7) It is desirable that SJS framework is accompanied with systems dynamic analysis of the interactions between each of the biophysical and social conditions. (8) Few dimensions related to PB framework that do not yet have any unanimously accepted indicators along with corresponding boundaries that fit national or sub-national scales, such as –change in biosphere integrity, stratospheric ozone depletion, ocean acidification, atmospheric aerosol loading and the introduction of novel entities. It is necessary. (9) Mere identification and measurement of indicators for the PB framework is not enough, even fail its purpose, if not proper checkpoints are made for these and implemented in policy. (10) Planning the future for nations under safe operating limits of biophysical resource consumptions and equitably provisioning social development should be the primacy in upcoming decades.

There are few novelties of this work: First, it aims to convey a pictorial portrayal of the dynamic state of socio-ecological indicators related to national-scale priorities and scenarios in the south and southeast Asia. Second, this work intersects a multidimensional set of indicators in a simple way, distinguishes the slit in the knowledge-base, and promotes new understandings to eliminate social deprivation whilst remaining under safe consumption limits of biophysical resources. Third, it provides south and southeast Asia’s (nation-scale) proximity to respective environmental boundaries and its satisfactory level of social wellbeing. Fourth, the sustainable development goals are “action-oriented, concise and easy to communicate, limited in number, aspirational, global in nature and universally applicable to all countries, while taking into account different national realities, capacities and levels of development and respecting national policies and priorities” (Rio+20 outcome document, 2012). Sixteen of the original seventeen SDG criteria have been connected in this framework. Thus, this work maintains a balance between simple and comprehensive approach, so that progress of all the SDGs in countries of south and southeast Asia can be comprehended (at least through 1 indicator/1 goal of UN SDG).

## Acknowledgement

We would like to thank Sk. Rohan Tanvir, The Institution of Engineers (India) for his assistance during the preparation of diagrams.

